# A Single-Cell Atlas of PVOD Across the Lifespan Reveals Cellular and Molecular Therapeutic Targets

**DOI:** 10.64898/2026.10.12.758636

**Authors:** Prashant Chandrasekaran, Sylvia N. Michki, Jennifer Pogoriler, Quinn Wright, MinQi Lu, Andrew R. Peterson, Hannah Hallquist, Edward Cantu, Edward E. Morrisey, Lisa R. Young, Maria C. Basil, Susan M. Lin, David B. Frank

## Abstract

**Rationale:** Pulmonary veno-occlusive disease (PVOD) is a progressive and lethal disease cured only lung transplant. While genetic mutations in *EIF2AK4* have been identified in some cases, 70% of PVOD patients have no known etiology.

**Objectives:** To construct a single-cell atlas to define the cellular and molecular constituents driving pathophysiology in PVOD.

**Methods:** Lung samples were collected from four idiopathic non-*EIF2AK4* mutation pediatric and adult PVOD patients and age-matched controls. Analyses of single-cell RNA sequencing, Xenium spatial transcriptomics, and immunohistochemistry were performed to identify cell state changes and differential gene expression between controls and PVOD and subsequent transcriptome comparison between our PVOD and published pulmonary arterial hypertension (PAH) datasets.

**Measurement and Main Results:** The single-cell atlas reveals marked changes in the PVOD endothelium compared to controls. EndMT and ectopic COL15A1+ systemic venous shunts and pulmonary vein endothelium are hallmarks of PVOD. Molecular analyses reveal activation of the integrated stress response (ISR)/ATF4, TGFβ signaling, MAPK signaling, and BRD4 are potential key players in the pathophysiology of PVOD. While PVOD and PAH share both pathological and some molecular mechanisms, ISR and TGFβ signaling are more prominent in PVOD than PAH.

**Conclusions:** These data indicate that endothelial dysfunction is a primary etiology in the development of PVOD with activation of multiple pathways with druggable targets.

**At a Glance Commentary:** *Scientific Knowledge of the Subject:* Pulmonary veno-occlusive disease (PVOD) is driven by progressive vasculopathy of the pulmonary veins and capillaries that results in significant pre-lung transplant mortality. Only 30% of PVOD cases can be explained by genetic mutations in *EIF2AK4*, a critical protein translation factor associated with the integrated stress response. Thus, the majority of PVOD cases have no known molecular and cellular etiologies.

*What This Study Adds to the Field:* This study generated a cell atlas of idiopathic PVOD in patients with the more common, non-*EIF2AK4* mutation etiology. This cell atlas defines pathological cell state changes in the lung vasculature including aberrant populations in the endothelium of veins and capillaries. These data also provide key insight into the molecular underpinnings driving PVOD disease and identified both novel and common targetable pulmonary vascular disease pathways with known drugs that are currently under investigation or FDA approved.

## Introduction

Pulmonary veno-occlusive disease (PVOD) is a rare but lethal disease (1–3). It is marked by venous endothelial cell (EC) sclerosis and dysfunction, neointima formation and capillary proliferation as found in pulmonary capillary hemangiomatosis (PCH). PVOD pathophysiology results in obliterative small vein and venule lesions leading to elevated pulmonary venous and arterial pressures, right heart failure, and death (3). Clinically, PVOD presents with rapidly progressive disease with high mortality and no curative treatments (3). While PVOD is estimated to cause 0.7-2.0% of all cases of pulmonary arterial hypertension (PAH) (4, 5) and shares similar pathology, the initiation of standard PAH therapy in PVOD can lead to life-threatening pulmonary edema. In the absence of pharmacological treatment options, lung transplant remains the only definitive treatment for PVOD.

PVOD pathogenesis involves complex interplay of endothelial dysfunction, aberrant vascular remodeling, and heritable genetic predisposition at cellular and molecular level. Recent studies uncovered mutations in *EIF2AK4* as a leading genetic cause in up to 30% of patients with PVOD (1, 2, 6, 7). EIF2AK4 or GCN2 is one of four kinases that incite the integrated stress response (ISR) that both attenuates protein translation and selectively upregulate translation of ATF4, a transcriptional regulator of adaptive responses to cellular stress (8). While recent data indicates maladaptive ISR contributes to chemotherapy-induced PVOD in rats (9, 10), loss of EIF2AK4 alone in animal models does not recapitulate the PVOD phenotype (6, 11, 12). Moreover, genetic mutations have been identified in only 30% of patients with PVOD, suggesting that unknown additional pathways may incite the pathophysiology of PVOD in most patients (7).

To identify potential cellular and molecular mediators of PVOD, we constructed a comprehensive cell atlas of idiopathic, mutation negative PVOD using single-cell RNA sequencing (scRNA-seq). We characterized novel cell states and transcriptomic changes in PVOD lungs and compared our findings to published PAH datasets (13, 14). Additionally, we confirmed and spatially mapped both RNA and protein level alterations in pulmonary vasculature by tissue immunofluorescence and spatial transcriptomics, respectively. Together, this study uncovers novel cellular and molecular mechanisms in the pathogenesis of PVOD with discovery of new potential therapeutic targets.

## Methods

Details on protocols for histology and methods used to assess both single-cell RNA sequencing and spatial transcriptomics can be found in the online data supplement.

### Human lung tissue sample collection

Unaffected human lung tissues were obtained from Human Lung Tissue Biobank in University of Pennsylvania Lung Biology institute. Briefly, these tissues were acquired from deceased donors with approved secondary use of tissues. Samples were screened negative for HIV and hepatitis A, B, and C. PVOD tissues were acquired for research purposes after informed consent. For both unaffected and diseased lung tissues, any information used to identify samples were removed in accordance with the Children’s Hospital of Philadelphia, Hospital of the University of Pennsylvania, and NIH protocols.

### BMP6 quantification

Volumetric analysis of BMP6 expression in unaffected and PVOD lung sections were quantified with IMARIS 10.0 (Bitplane) using previously established procedures (15). Briefly, from the lung section, five regions were selected for each tissue. Using ‘volume statistics’ function in IMARIS, total lung volume was obtained from surface function of DAPI channel. To measure the expression of BMP6, single channel of BMP6 was reconstructed in 3D using the surface function. The density was then calculated by dividing BMP6 volume by total lung volume.

### Phospho-SMADs, BRD4, ATF4 and EIF2AK2 quantifications

Quantifications were done without segregating capillaries and microvessels. Briefly, five regions covering capillaries and microvessels, and only veins were imaged on both unaffected and PVOD tissues. In case of capillaries and microvessels, a pan endothelial marker ERG was used to identify the endothelial cells. Using ‘colocalization function’ in IMARIS, endothelial cells expressing phospho-SMADS (pSMADs), BRD4, ATF4, and EIF2AK2 were isolated. ‘Spots function’ was then used to count the number of cells in each instance. We identified the percentage of cells expressing the relevant marker by dividing colocalized cells with ERG+ cells. FIJI was used to count the cells expressing pSMADs, ATF4, and EIF2AK2 in the veins.

### Statistics

Unpaired 2-tailed *t*-tests were used to detect significant differences in quantification analyses. All statistical tests were performed using Prism 10.0, with significance detected at: \**P* ≤ 0.05. Each point on the graph denotes a biological repeat and specific *n* values have been reported in figure legends.

## Results

### A Single-Cell Atlas Identifies Pathological Cellular Heterogeneity Underlying PVOD

To identify pathological cell states and molecular mechanisms underlying PVOD, we obtained lung tissue from the Penn-CHOP Lung Biology Institute Human Lung Tissue Biobank that included explanted lungs of four pathology-confirmed PVOD patients and three age-matched control patients (age range of 12 years to 37 years old; Table 1). We confirmed the classical phenotype in PVOD using hematoxylin and eosin (H&E) staining, identifying luminal occlusion with neointima and intimal sclerosis and significant muscularization of microvessels and veins in affected tissues (Figure 1A and Figure E1A-D). To identify the cellular and transcriptomic changes in PVOD, we prepared single cell suspensions from whole lung tissue and sorted cells using a magnetic beads-based approach to enrich for CD45-cell populations. Two samples from each patient were collected and sequenced including a CD45-cell sample and CD45+ cell sample. After filtering out dead and low-quality cells using quality assessment metrics, batch correction and dimensionality reduction, we profiled a total of 38,917 cells from PVOD and unaffected samples. Unbiased Leiden clustering of integrated dataset revealed 29 transcriptomically distinct cell clusters, and 26 different cell types with high fidelity (Figure 1B-D and Figure E1E). We compared cell proportions between PVOD and unaffected and revealed significant shifts within both mesenchymal and endothelial compartment. Interestingly, PVOD tissues showed marked expansion of vascular smooth muscle cells (VSMCs) and pericytes within the mesenchyme (Figure 1E). Additionally, there was a significant expansion of general capillaries (gCap or CAP1) within the endothelial compartment reminiscent of capillary proliferation seen in PVOD and PCH (Figure 1F).

**Figure 1.**
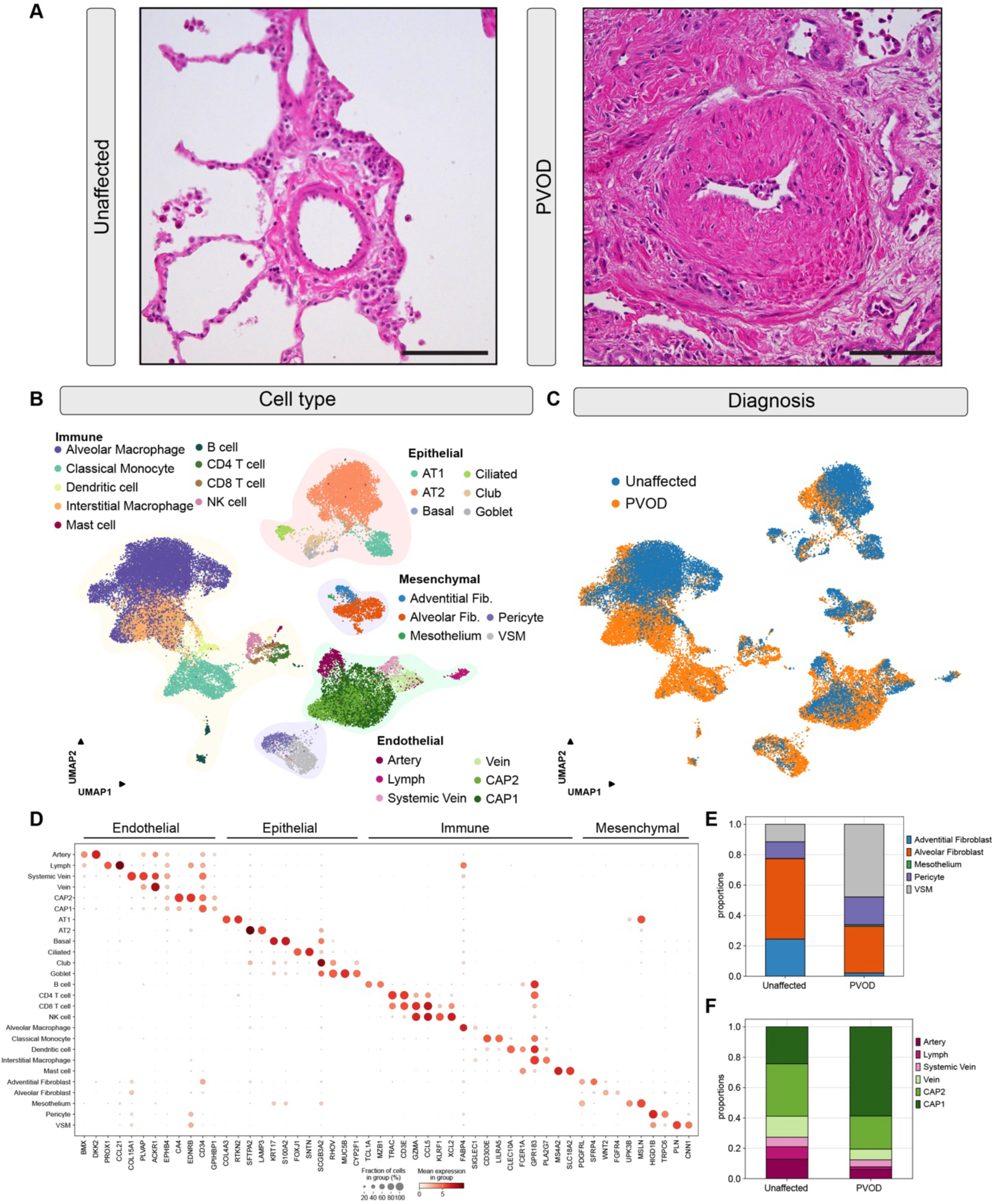
A Single-Cell Atlas of PVOD. (A) Images of H&E-stained tissue from unaffected (left) and PVOD (right) lungs, showing neointimal thickening in PVOD affected vein. (B) Uniform Manifold Approximation and Projection (UMAP) of scRNA-seq data with cells from unaffected and PVOD lungs colored by cell type. (C) UMAP embedded with cells colored by diagnosis (unaffected/PVOD). (D) Dotplot showing expression and fraction of selected cell type marker genes across all cell clusters. (E) Stacked barplot showing the relative proportions of mesenchymal cells in unaffected and PVOD lung derived scRNA-seq data. (F) As in (E), for endothelial cells. Scale bar: 100µm.

**Table 1.** Characteristics of PVOD patients in cell atlas.

| Table 1. Characteristics of PVOD patients in cell atlas |  |  |  |  |
| --- | --- | --- | --- | --- |
| Patient | Age | Gender | WES | PVOD subtype |
| 1 | 12 | Male | Negative | Idiopathic |
| 2 | 15 | Male | MYBPC3 | Idiopathic |
| 3 | 31 | Female | Negative | Granulomatous |
| 4 | 37 | Male | Negative | Idiopathic |
| 5 | 12 | Female | N//A | Control |
| 6 | 17 | Male | N/A | Control |
| 7 | 35 | Female | N/A | Control |

### PVOD is Associated with an Endothelial-To-Mesenchymal Transitional (EndMT) Cell State

EndMT is critical for not only normal vascular development, but it also underlies pathology in many cardiovascular diseases including pulmonary hypertension and PVOD (16–19). Additionally, while possibly not a direct contributor to neointima, EndMT likely plays an indirect role in guiding neointima formation (20). To assess the gene related changes specific to endothelium and EndMT and to unravel the interactions with mesenchyme, we re-clustered the endothelial and mesenchymal populations from our dataset, identifying 10 transcriptomically distinct cell types (Figure 2A). On assessment of the EC marker, *VWF* and mesenchymal marker *TAGLN*, we noted co-expression in multiple EC populations (Figure 2B-D). Notably, the largest population with overlapping expression of these two markers was the pulmonary vein EC population, consistent with an EndMT state. To corroborate this state change, dot plots for expression of EndMT related genes in pulmonary vein ECs including multiple EndMT-related transcription factors and the TGFβ signaling reporter gene, *SERPINE1*, were generated (Figure E2A). We observed significant activation of TGFβ signaling and upregulation of *ZEB1*, supporting an EndMT state. We further characterized these cells spatially using immunohistochemistry (IHC) by staining for VWF and TAGLN and found them distributed in endothelial/intimal in PVOD samples only (Figure 2E and 2F).

**Figure 2.**
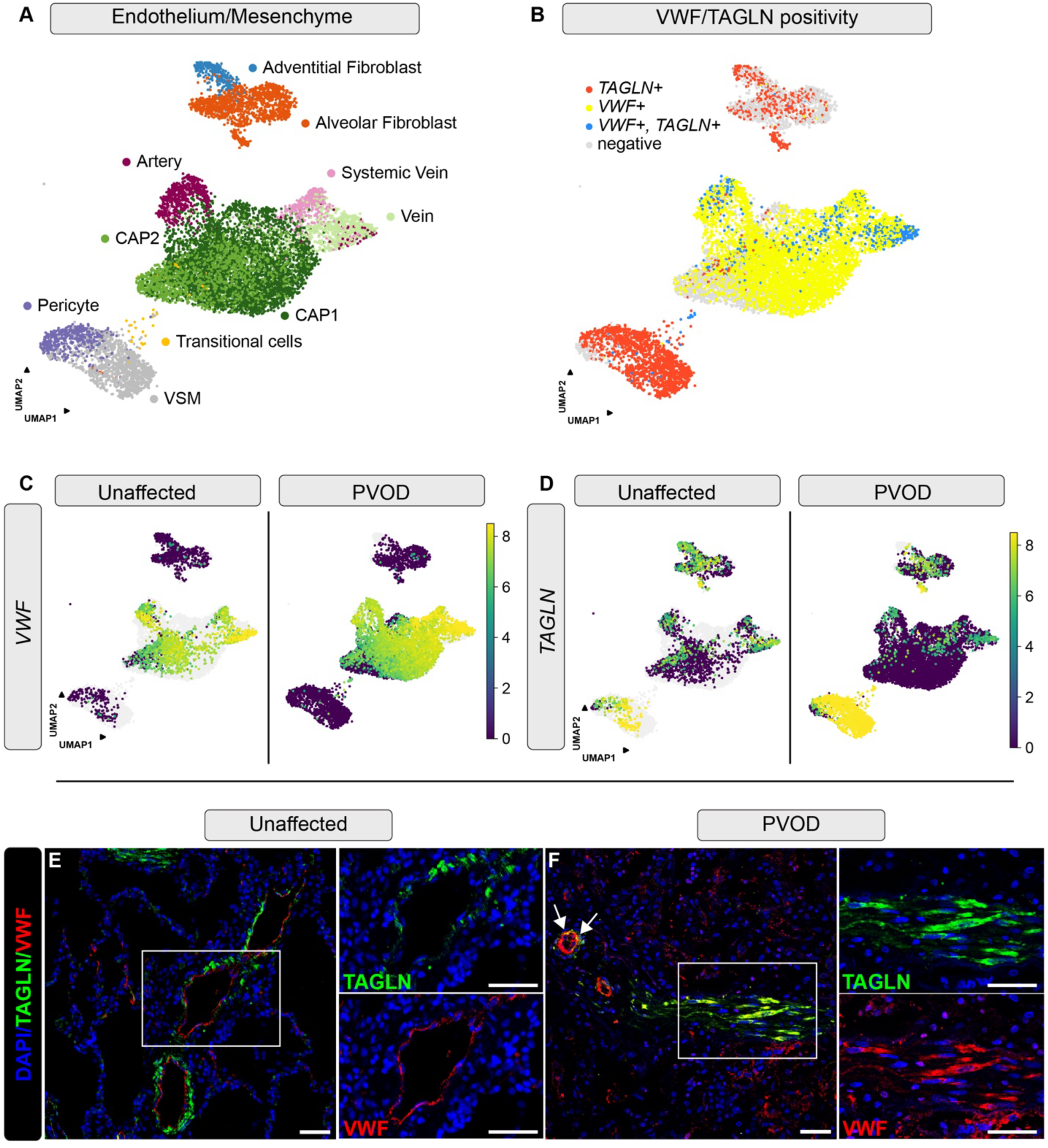
Endothelial-to-mesenchymal (EndMT) changes in vein endothelium. (A) UMAP of embedded endothelial and mesenchymal cells colored by cell type. (B) UMAP of embedded endothelial and mesenchymal cells colored by *TAGLN+*, *VWF+*, or *TAGLN+/VWF+* (greater than 1 UMI for transcript of interest). Cells with no *TAGLN* or *VWF* UMIs detected are classified as negative. (C-D) UMAPs of embedded cells colored by *VWF* (C) and *TAGLN* (D) gene expression and split by condition. (E-F) IHC staining for TAGLN, VWF and DAPI in unaffected (E) and PVOD (F) lung tissue. Boxed areas are magnified and depicted to the right with TAGLN or VWF only. (F) White arrows point to positive cells in small vessels and boxed area represents a longitudinal section of vessel. Scale bars: 50µm.

### COL15A1 Marks Ectopic Lung Parenchymal Shunts and EC Remodeling in PVOD

We further dissected the endothelial compartment and identified previously described EC populations including the two transcriptomically distinct vein EC clusters, namely systemic vein marked by expression of *COL15A1* and pulmonary vein cluster marked by expression of *ACKR1* (Figure 3A) (21). COL15A1 is a non-fibrillar collagen associated with basement membranes, and its importance has also been studied in physiological and pathological responses such as atherosclerosis (22, 23), and more recently in the lung as ectopic shunt vessels observed in alveolar capillary dysplasia (ACD), pulmonary fibrosis, and lung cancer (24–28). Given these lung pathologies, we asked how *COL15A1* was altered in PVOD. UMAPs of combined unaffected and PVOD cells colored by expression of *ACKR1* and *COL15A1* demonstrated expansion of *COL15A1* into pulmonary vein ECs and CAP1s (Figure 3B-D). We observed single *ACKR1*+ pulmonary vein ECs, double *ACKR1*+/*COL15A1*+ systemic vein ECs, and single *COL15A*1+ cells (Figure 3B). UMAPs of separated unaffected and PVOD cells colored by individual expression of *COL15A1* (Figure 3C) or *ACKR1* (Figure 3D) revealed that this expansion of double *ACKR1*+/*COL15A1*+ systemic vein ECs and single *COL15A1*+ cells were largely found in PVOD samples. To identify these cell types spatially, we performed IHC for COL15A1 and ACKR1 proteins (Figure 3E-F and E3A-C). While systemic vein ECs were present in vessels surrounding large airways of both unaffected and PVOD samples, only PVOD samples exhibited double ACKR1+/COL15A1+ shunt vessels and single COL15A1+ vessels in the distal lung parenchyma.

**Figure 3.**
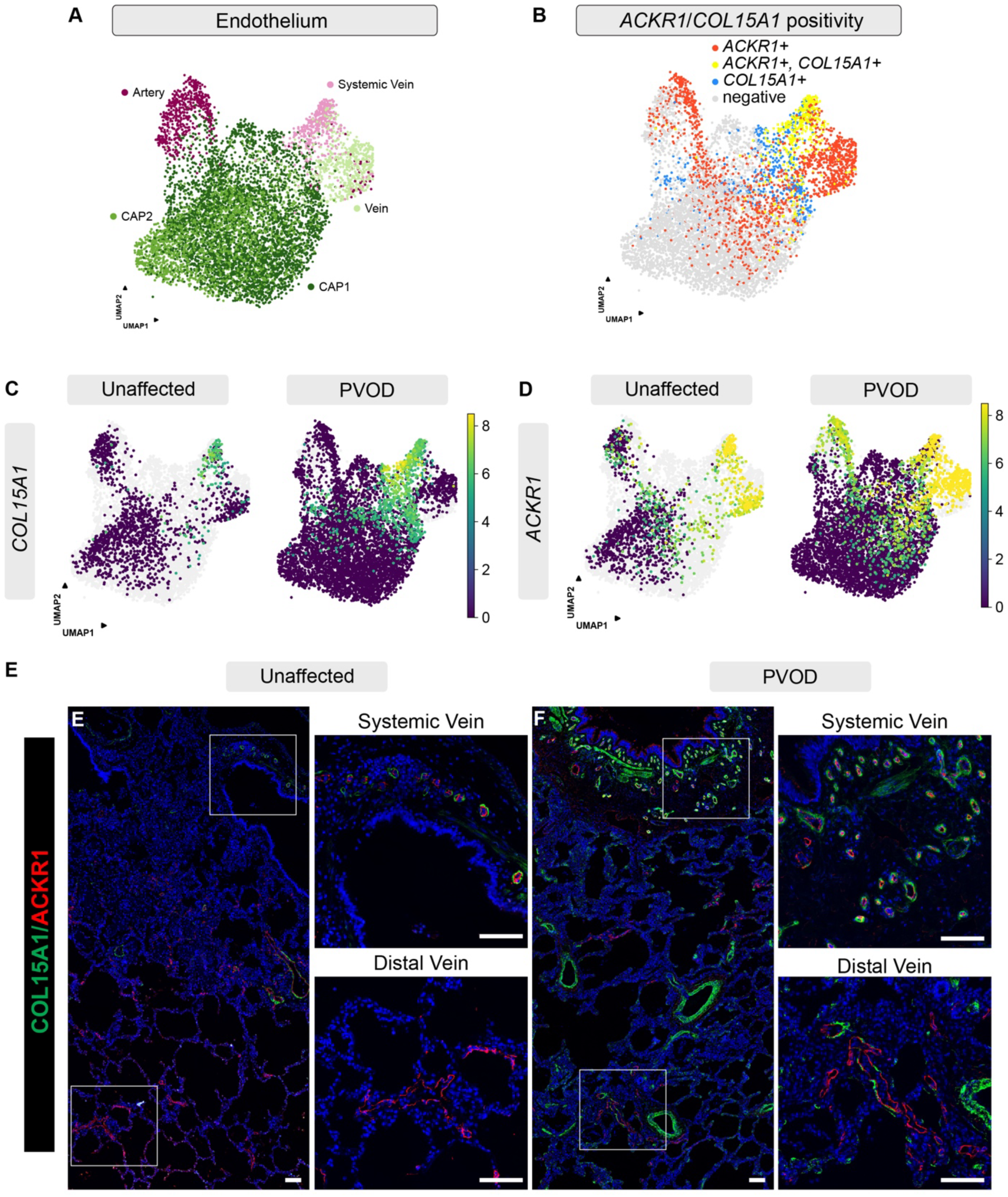
Expansion and ectopic expression of systemic vein into lung parenchyma in PVOD. (A) UMAP of embedded ECs colored by EC subtype. (B) UMAP of embedded ECs colored by expression of *ACKR1+*, *COL15A1+*, and *ACKR1+/COL15A1+* (greater than 1 UMI for transcript of interest). Cells with no *ACKR1* or *COL15A1* UMIs detected are classified as negative. (C-D) UMAP of embedded ECs colored by *COL15A1* (C) and *ACKR1* (D) gene expression split by conditions. (E-F) IHC staining for ACKR1, COL15A1and DAPI in unaffected (E) and PVOD (F) lung tissue, split by systemic veins and distal veins. Scale bars: 100µm.

### Dysregulation of ISR is Associated with Maladaptive and Increased ATF4 Expression

We continued to focus on the endothelium and generated UMAPs embedded with ECs colored by cell subtype (Figure 4A). We then performed Overrepresentation Analysis (ORA) of Gene Ontology (GO) categories of differentially expressed genes (DEGs) to predict biological functional differences between PVOD and unaffected tissues and observed enrichment of stress-related response pathways (eukaryotic translation initiation factor 4F) and pathways related to cell adhesion and spreading, consistent with activation of ISR (Figure 4A) (9, 10, 29–33). We focused on components of the ISR including Activating transcription factor 4 (ATF4) and protein translation factor, EIF2A, and its associated kinases and feedback molecules. Additionally, we examined a differentially expressed regulator of ATF4, Bromodomain protein containing 4 (BRD4), and a downstream mediator of ISR, endothelin-1 (EDN1) (34, 35). We generated dot plots and UMAPs to assess expression of these components (Figure 4B and E4A-D). While we observed no changes in enrichment of EIF2AK4, there were significant increases in expression of *ATF4*, *EIF2AK2*, and *GADD45A*. Moreover, both *BRD4* and *EDN1* were highly enriched in PVOD endothelium. To validate these findings, we performed IHC for ATF4, EIF2AK2 (PKR), and BRD4 in combination with pan-endothelial marker ERG and quantified their expression in endothelium and noted a significant increase in the number of ATF4, EIF2AK2, and BRD4 ECs (Figure 4C-K). Recently, macrophage ferroptosis and venous arterialization has been implicated in the pathogenesis of PVOD related to *EIF2AK4* mutations (33). Using markers of ferroptosis identified through the ferroptosis database, FerrDb V3 (36), we generated heatmap of markers of ferroptosis in monocytes/macrophages (Figure E4E). We observed no significant changes in ferroptosis markers other than *ACSL4*, and the critical regulators of ferroptosis, *HMOX1* and *TFRC* were found decreased in PVOD compared to unaffected controls (Figure E4F-I).

**Figure 4.**
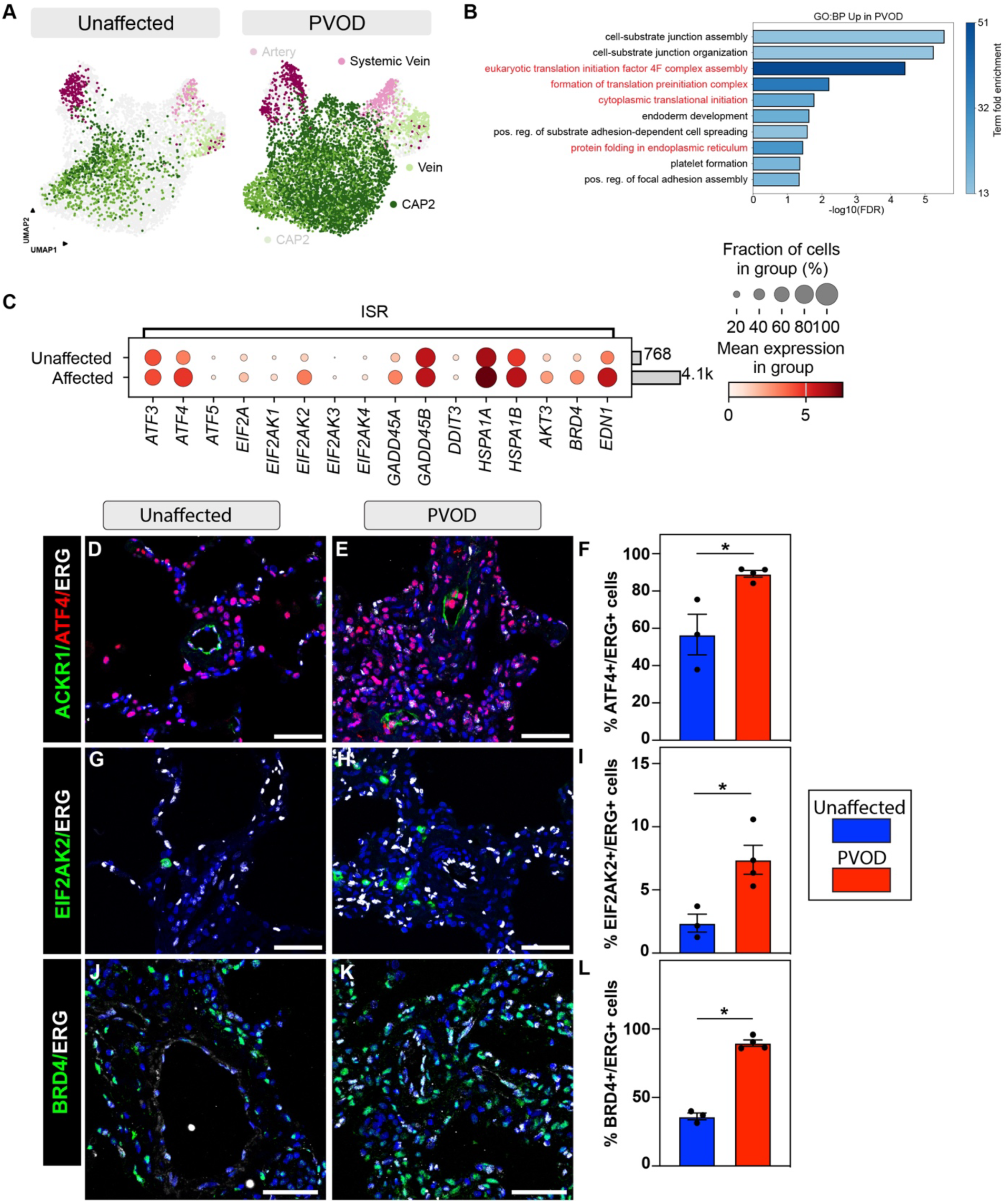
Increased ISR pathway response genes in PVOD. (A) UMAP of embedded ECs colored by EC subtype, split by unaffected (left) and PVOD (right) status, with cell types of interest highlighted. (B) Overrepresentation analysis (ORA) of Gene Ontology (GO) terms of genes in PVOD compared to unaffected. (C) Dotplot showing expression of selected ISR pathway related cell type marker genes across all ECs. (D-L) IHC staining for ATF4, ACKR1 and ERG (D, E), EIF2AK2 and ERG (G, H), and BRD4 and ERG (J, K). Quantification of percentage of ATF4+ (F), EIF2AK2+ (I), and BRD4+ (L) ECs, (n = 3 for unaffected, n = 4 for PVOD, data represented as mean ± SEM, * *P* ≤ 0.05). Scale bars: 50µm.

### MAPK and TGFβ pathways are Active in PVOD Endothelium

Given the presence of EndoMT cells and expansion of COL15A1+ cells in the CAP1s and vein clusters, we probed for additional molecular pathways that contributed to the underlying pathology (Figure 5A). We assessed differentially expressed genes in PVOD tissues and identified marked increase in *MAP2K2*, a core kinase of MAPK/ERK signaling cascade (Figure 5B-E). Using weighted pathway analyses, we observed that PVOD tissues exhibited increases in MAPK, hypoxia, and JAK-STAT pathway activity (Figure 5C). We previously noted increased *SERPINE1* expression indicating increased TGFβ activity in PVOD (Figure E2A). TGFβ signaling is a critical and common pathway in pulmonary vascular disease (37). To confirm TGFβ activity in the endothelium, we first generated dot plots of differentially regulated TGFβ signaling genes and noted a significant enrichment in signaling components including *BMP6*, *TGFBR2*, and *TGFB1* (Figure 5D and E5A-J). Additionally, we used the pathway analysis described above to plot expression of TGFβ pathway activity across the endothelium, PVOD samples showed higher TGF-β activity (Figure 5E). Next, we performed IHC for BMP6 and phospho-SMAD1/5/9 and phospho-SMAD2 to measure BMP and TGFβ signaling activation, respectively. In combination with pan-endothelial marker ERG, we found a trend in increase in expression of BMP6 and phospho-SMAD2+ ECs in the PVOD endothelium but noted no change in BMP signaling.

**Figure 5.**
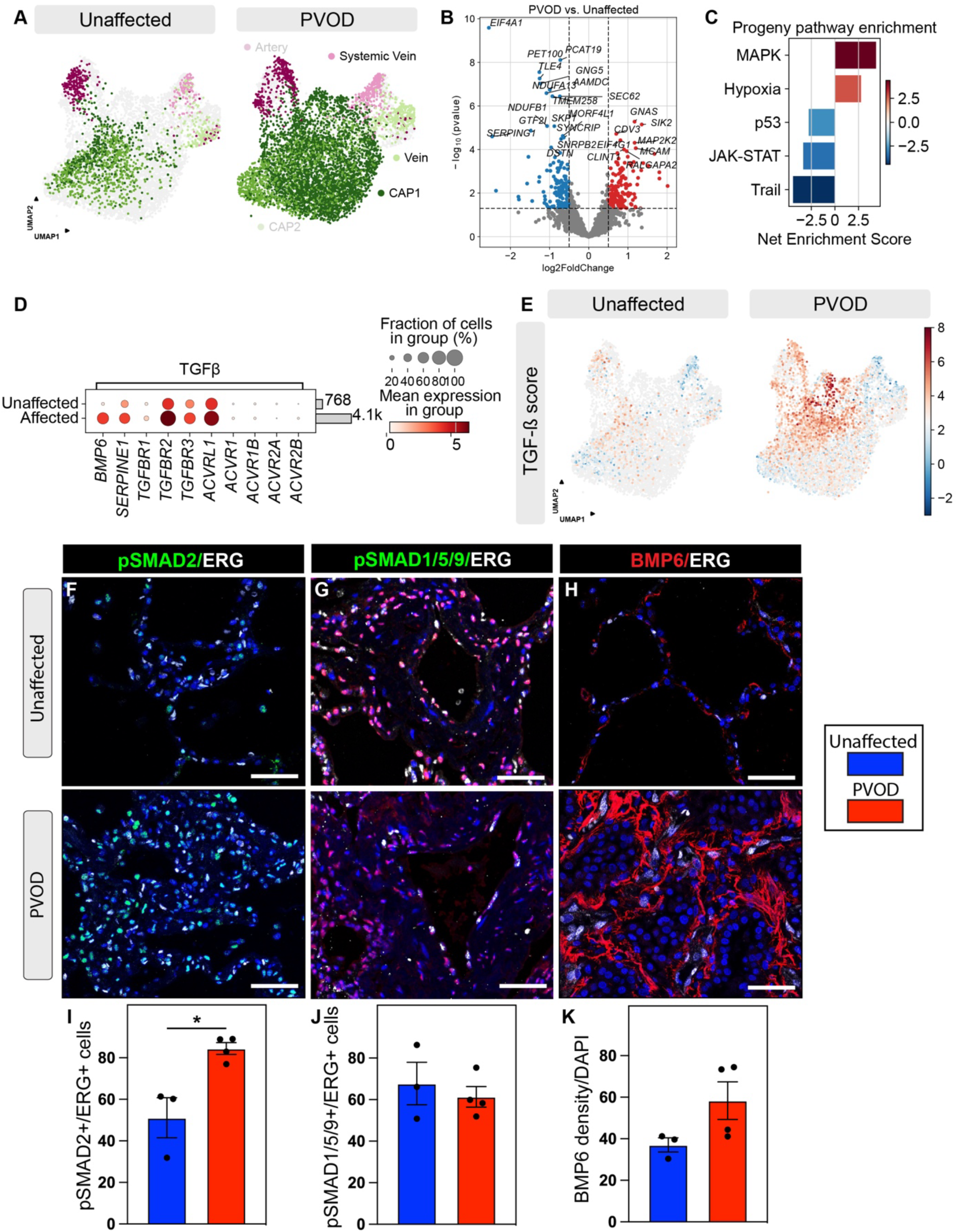
Increased MAP kinase and TGF-ß pathway activity in the endothelium in PVOD. (A) UMAP of embedded cells colored by EC subtype and split by unaffected (left) and PVOD (right) status. (B) Volcano plot of pseudobulk differential expression testing results for Systemic Vein, Vein, and gCap/CAP1 cells in PVOD vs. unaffected controls. (C) Enrichment scores for pathways in the Progeny pathway database based on differentially expressed genes in (B). (D) Dotplot showing expression of selected TGF-ß pathway related cell type marker genes across all cells in the scRNA-seq atlas, split by conditions. (E) UMAP showing TGFβ Progeny pathway enrichment scores from (C) in unaffected (left) and PVOD (right) endothelial cells. (F-K) IHC staining for ERG and phospho-SMAD2 (pSMAD2) (F), phospho-SMAD1/5/9 (G), and BMP6 (H) in unaffected (top panels) and PVOD (bottom panels) tissue. Quantification of number of cells positive for pSMAD2 expression (I) and pSMAD1/5/9 expression (J) and density of BMP6 expression (K), (n = 3 for unaffected, n = 4 for PVOD, data represented as mean ± SEM, *: *P* ≤ 0.05). Scale bars: 50µm.

To further refine spatially distinct signaling pathways, we performed spatial transcriptomics using 10X Genomics Xenium 5K that allows for the detection of 5,000 human genes. PVOD pathology was confirmed using H&E (Figure 6A). Using K-mean-based clustering, we defined the major cell compartments including endothelium, mesenchyme, epithelium, and immune cells (Figure 6B). We next focused on identifying pulmonary veins that were large unaffected (Figure 6C-F) and veins that had undergone extensive remodeling and neointima formation (Figure 6G-J). Vein and artery ECs were defined by the previously reported gene markers including *EPHB4*, *ACKR3*, and *SELP* and the markers *GJA5*, *DLL4*, and *SOX17*, respectively (21). VSMCs and previously described NOTCH3+ VSMCs of the neointima were marked by *NOTCH3*, *PDGFRB*, and *ACTN1*, which were increased in veins with extensive remodeling (Figure 6J) (20). Additionally, many of the veins with significant remodeling and neointima expresses some transcripts specific to artery ECs, suggesting arterialization of some veins (Figures 6F and 6J). Given the presence of TGFβ activation in PVOD, we examined the expression of *SERPINE1* and observed a significant increase in transcripts in extensively remodeled veins (Figures 6F and 6J). We further defined the TGFβ signaling by mapping the expression of previously identified differentially regulated genes in addition to the activin signaling branch of the TGFβ superfamily. Both TGFβ ligands and receptors are expressed in ECs and mesenchyme (Figure E6A and E6B). Interestingly, while the activin B ligand, *INHBB*, is expressed in the endothelium and epithelium, the activin A ligand, *INHBA*, is almost exclusively found in macrophages (Figure E6C). However, receptors for activin signaling are expressed in most cell compartments (Figure E6D).

**Figure 6.**
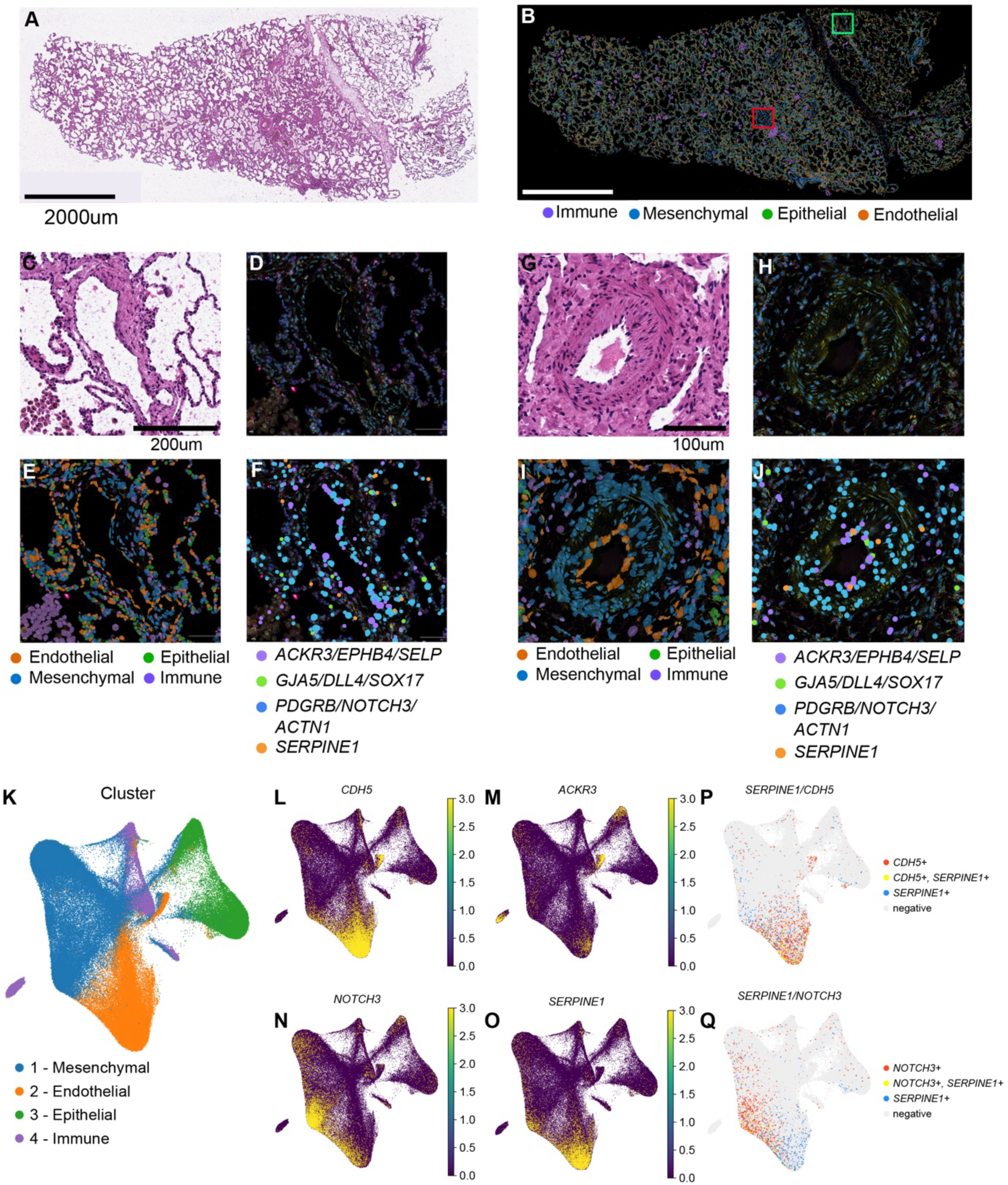
Spatial transcriptomic analysis of PVOD tissue defines differential TGF-ß activity in unaffected vein vs affected vein. (A) H&E staining of PVOD tissue. (B) PVOD tissue referenced in (A), colored by immune, mesenchymal, epithelial and endothelial cluster assignments. (C) H&E staining of unaffected vein. (D) Xenium morphology staining of unaffected vein in (C). (E) Cell segmentation boundaries colored by cell type in (B). (F) Transcripts for arterial (green), venous (purple), mesenchymal (blue), and the TGFβ responsive gene *SERPINE1* (orange) in unaffected vein. (G) H&E staining of affected vein. (H) Xenium morphology staining of unaffected vein in (G). (I) Cell segmentation boundaries colored by cell type in (B). (J) Transcripts for arterial (green), venous (purple), mesenchymal (blue), and the TGFβ responsive gene *SERPINE1* (orange) in affected vein. (K) UMAP obtained from xenium data colored by cell type. (L-O) UMAP of all cells colored by gene expression of *CDH5* (L), *ACKR3* (M), *NOTCH3* (N) and *SERPINE1* (O). (P) UMAP of all cells colored by *CDH5+*, *SERPINE1+*, and *CDH5+/SERPINE1+* and Q) UMAP of all cells colored by *NOTCH3+*, *SERPINE1+*, and *NOTCH3+/SERPINE1+*. Scale Bar: A-B=2000µm, C-F=200µm, and G-J=100µm.

### PVOD and PAH are marked by distinct cellular and molecular dysfunction

Given similarities in pathology and mechanisms, we sought to delineate the underlying differences in gene dysregulation informing cellular processes distinguishing PVOD from PAH. We integrated our PVOD dataset with PAH and control datasets from Kropski et al. 2025, Tuhy et al. 2025 and Valzano et al. 2022 (GSE296823, GSE293580 and GSE210248 respectively). Unbiased Leiden clustering identified 26 transcriptomically distinct cell types (Figure 7A-B), wherein we resolved differential activity within endothelial clusters of Vein, Systemic Vein and CAP1s. Examining the expression of core cellular components involved in ISR and TGF-β signaling cascades, we found marked upregulation of ISR related genes like *ATF4*, *EIF2AK2*, *GADD45A*, *GADD45B*, *HSPA1A* and *HSPA1B*, alongside TGF-β/EMT associated genes including *BMP6*, *SERPINE1*, *TGFBR2* and *HES1* in PVOD endothelial clusters (Figure 7B). We further performed differential gene expression testing within VSMC cluster across PAH and PVOD. Relative to PVOD, VSMCs of PAH tissues showed expression of *CD74*, *NUPR1* and *PDFN5* (Figure 7C), whereas in PVOD, VSMCs showed increased expression of *JUN*, *HSPA1A*, *ATF3* and *HIF1A*, among others (Figure 7C). Further, PROGENy pathway activity analysis in the VSMC cluster identified increased MAPK, PI3K, and JAK-STAT pathway activities, once-again highlighting the preferential activation of MAPK signaling in PVOD relative to PAH (Figure 7D). We also performed ORA on DEGs to predict biological functional differences between PVOD and PAH tissues and predicted enrichment of integrated stress response signaling, responses to protein refolding and host-mediated activation of viral transcription (Figure 7E).

**Figure 7.**
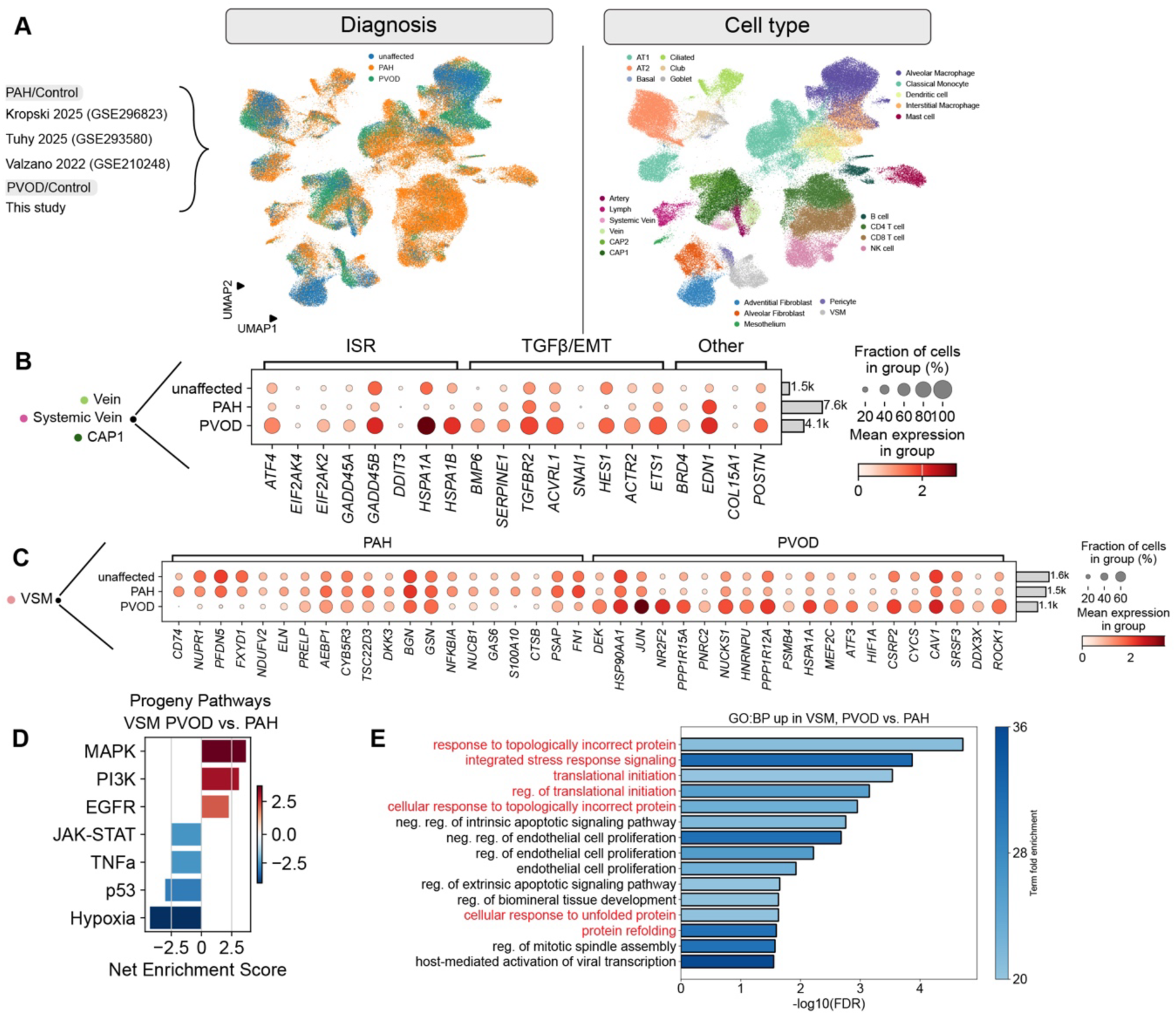
A comparison of differential gene regulation in PVOD versus pulmonary arterial hypertension (PAH) highlights integrated stress response and TGFβ activity in endothelial and mesenchymal cells. (A) UMAP overview of the integrated scRNA-seq dataset, colored by diagnosis (left) and cell type (right). Datasets included in this integration are listed on the far left. (B) Dotplot of expression of selected integrated stress response (ISR), TGFβ/EndMT, and other genes of interest in Systemic Vein, Vein, and gCap cells. (C) Dotplot of expression of genes differentially expressed in vascular smooth muscle (VSM) cells: a PAH vs. PVOD comparison. Expression in unaffected cells is plotted alongside the tested conditions. (D) Enrichment scores for pathways in the Progeny pathway database based on DEGs identified in (C). (E) ORA results showing GO:BPs of DEGs up-regulated in PVOD from (C).

## Discussion

The pathogenesis of PVOD is dependent on a complex interplay of endothelial dysfunction, aberrant vascular remodeling, and heritable genetic predisposition. In the absence of any effective therapeutics, PVOD is lethal leaving lung transplant as the only option. We collected a cohort of idiopathic, *EIF2AK4* mutation-independent PVOD samples and found abnormal venous and capillary cell state changes defined by EndMT, COL15A1+ venous ECs, and potential venous arterialization. Additionally, we observed maladaptive ISR and TGFβ superfamily defects consistent with previous studies examining both rodent and human PVOD (9, 10, 17, 29, 31, 32). Moreover, we have defined several novel pathways with druggable targets, uncovering potential disease-specific therapies for PVOD.

Our data reveal a novel population of ACKR1+COL15A1+ cells in the distal vasculature. Reflecting anatomical differences in pulmonary and bronchial circulation, COL15A1 is normally restricted to systemic veins (21). As expected, we found COL15A1+ systemic vein ECs surrounding large airways in both control and PVOD samples. Beyond the systemic veins, we also found ectopic expansion of COL15A1 into the distal vasculature with ACKR1+/COL15A1+ shunt vessels and COL15A1+ veins in PVOD samples. We subsequently confirmed the expansion and displacement of COL15A1 in distal veins and gCap/CAP1s by immunofluorescence. This phenomenon has been observed in other pulmonary diseases. The replacement of maldeveloped capillaries by COL15A1+ vessels was recently described in ACD and interstitial lung diseases (25, 26, 38). Moreover, in chronic thromboembolic pulmonary hypertension, expansion of the COL15A1+ cell population was shown to be with recanalization. This paradigm has recently expanded to PAH, in particular to systemic scleroderma associated PH (SSc-PH) (39). In PVOD, these COL15A1+ cells likely represent neovascularization of venous ECs with subsequent shunting through malformed pathways. To confirm this will require advanced 3D imaging of thick tissue sections to fully characterize whether COL15A1+ and COL15A1+/ACKR1+ endothelium represent de novo systemic venous shunts versus cell state changes involving remodeling of basement membrane and stiffening of existing pulmonary veins (22, 23, 40).

While the primary affected endothelial cell populations differ, similar clinical and histological features exist between PVOD and other forms of PAH, and it has long been suspected that multiple forms of PAH and PVOD have shared molecular pathways (31, 32). EndMT and its related gene signature including TGFβ signaling are common mechanisms shared by both PVOD and PAH. PAH is often defined by an imbalance in a two-pronged signaling system with pro-remodeling and EndMT dictated by TGFβ and activin signaling on one end and an opposing anti-remodeling branch guided by BMP signaling (41–43). This concept forms the cornerstone for the recently approved activin ligand trap, Sotatercept (44–46). Our data indicate a more prominent role for TGFβ signaling in PVOD versus PAH with gene enrichment in the TGFβ receptors and the target gene, *SERPINE1*. Interestingly, we see an increased in *BMP6* mRNA and trend in increase in BMP6 that binds to not only BMP receptors but also activin type 2 receptors and the ACTRII-Fc ligand trap (47, 48). Recent evidence points to a pathological role for BMP6 in vascular disease with a role in pathogenic angiogenesis and vascular calcification (49, 50). These findings provide evidence for the expanded use of ligand traps such as Sotatercept for the treatment of PVOD, which has recently been demonstrated efficacious in a several patients in a case study (16).

Our comprehensive cell atlas of idiopathic PVOD highlights the role of mammalian ISR in pulmonary vascular endothelial cell remodeling despite the absence of *EIF2AK4* driver mutation in our samples. Despite this, we see increased ATF4 in pulmonary vein and general capillary endothelium in vivo, suggesting not only an EIF2AK4-independent signaling mechanism for ISR in PVOD compared to previous studies but also highlights the importance of this pathway even in non-genetic causes of PVOD (31, 32). Proposed alternative pathways may include activation of EIF2AK2 (PKR) that we observed and has been validated in rodent models of PVOD (9, 10, 29).

There are several limitations to our work. Although we found ectopic lung parenchymal shunts and endothelial remodeling in PVOD, the specific role of COL15A1+ endothelial cells in shunts and neovascularization remains unclear and will require future collection of PVOD tissue to analyze using advanced 3D imaging. In addition, we found ACKR1+ cells in the arterial EC cluster which may represent arterialization of venous ECs. More work is needed to understand the implications of this cell population. We are further limited in that scRNA-seq examines expression of mRNA, and some of these components are regulated at the protein level. We have done diligence to look at many of these molecules at the protein by IHC and observed similar dysregulation for most molecules. Additionally, we focused on the endothelium and VSMC for most of this study, and there may be dysregulation in other niches not explored in detail here such as macrophages that have been implicated in *EIF2AK4* mutation patient and rodent samples (33). Our cell atlas is limited in that it does not include samples from patients with *EIF2AK4* mutations. However, as *EIF2AK4* mutations are found in only up to 30% of patients, and the mechanisms proposed by our cell atlas may represent the broader PVOD patient population (1, 2, 6, 7).

## Conclusion

In summary, we have constructed a comprehensive cell atlas of PVOD. We have defined cell state changes that may contribute to the pathophysiology of PVOD. Importantly, we have corroborated findings from foundational previous studies describing the role of ISR, TGFβ superfamily, and MAP kinase signaling (9, 10, 17, 29, 31, 32). Together, these data provide rationale for the expanded use of drugs that are FDA approved or in clinical trials including inhibitors to BRD4, the ISR, and MAPK/ERK, and TGFβ superfamily ligand traps such as Sotatercept. Once a lethal disease, previous studies and this work may provide therapies to reduce progression and improve outcomes for PVOD.

## Author’s contributions to the study

SML, MCB, EEM, LYR, and DBF conceived the project. PC, SML, SNM, MCB, and DBF designed the experiments. PC, SML, SNM, JP, QW, ML, ARP, HH, EC, EEM, LYR, MCB, and DBF conducted experiments and/or contributed to data analysis and interpretation. PC, SML, SNM, EEM, LYR, MCB, and DBF contributed to the writing and editing of the manuscript.

## Acknowledgements

We are forever thankful to the patients and families who have so graciously and selflessly contributed to this study. We would also like the thank Dr. Florin Tuloc and CHOP Flow Cytometry core for aiding in cell isolation and Dr. Dan Martinez and CHOP Pathology core and Dr. Mei Zhange and the CHOP Single-Cell Technology core for assistance with the Xenium platform.

## Data Availability Statement

Single-cell RNA sequencing data in this study have been deposited into Zenodo with DOI, https://doi.org/10.5281/zenodo.23166320.

