## Supplemental Methods & Figures for "A Single-Cell Atlas of PVOD Across the Lifespan Reveals Cellular and Molecular Therapeutic Targets"

**Supplementary Data Online**

**Methods**

**Single-cell RNA-Seq sample and Xenium sample preparation.** Single cell suspension of lung tissues was prepared based on previously established protocols (1). Briefly, unfixed human lung specimens were isolated by pathologist through direct visualization. These isolated specimens were minced using mechanical agitation and enzymatically digested using a cocktail of collagenase (Gibco, 17100017), dispase (Corning, 354235) and DNase I (ThermoFisher, EN0521) for 20 minutes at 37°C. Post digestion, samples were filtered using a 40μm filter (info), treated with ACK lysis buffer before performing bead-based sorting strategy. We selectively isolated $CD45^{+ve}$ cells using CD45 microbeads (Miltenyl Biotec, 130-045-801) and MACS LS column. Both $CD45^{+ve}$ and $CD45^{-ve}$ cells were concurrently loaded at a volume of 16,500 cells per each individual lane of a 10X Genomics 3' gene expression scRNA-seq assay chip G (v3.1, 10X Genomics) to recover 10,000 live cells. Further, region of interest from paraffin embedded lung tissues were visually identified by clinical pathologist and was isolated and sectioned on to a Xenium slide and processed further based on manufacture's protocol (10X Genomics).

**scRNA-Seq analysis and Xenium analysis.** scRNA-seq data generated/analyzed in this study were processed as previously published (2-7) and as follows. Reads in FASTQ format were aligned to the NCBI RefSeq GRCh38 genome and reads with unique molecular identifiers (UMIs) were quantified using STAR-solo, with *soloUMIdedup="1MM_CR"*, *soloUMIfiltering="MultiGeneUMI_CR",* and *soloCellFilter="EmptyDrops_CR"* (8)Ambient RNA was removed from counts matrices using the scvi-tools implementation of scAR (9, 10). Counts matrices were processed using the scanpy and anndata python libraries (11). Cells were filtered to include only those cells with more than 500 unique genes detected/cell, fewer than 6500 unique genes detected/cell, and fewer than 10% of UMIs attributed to mitochondrial genes. Genes were excluded if detected in fewer than 10 cells. Putative cell doublets were identified and removed from each batch independently using the scvi-tools implementation of the SOLO doublet identification model (12).

Gene expression was calculated from scAR-corrected UMI-counts by first normalizing to counts per 10K UMIs (CP10k) using the *sc.pp.normalize_total* function with *target_sum=10000* and then calculating ln(CP10k + 1) using the *sc.pp.log1p* function. The top 2000 highly variable genes in the dataset were identified using the *sc.pp.highly_variable_genes* function, with *flavor="seurat_v3"* and *n_top_genes=2000*. A latent representation of the data was generated using scVI (13), with *n_genes_by_counts* included as a continuous covariate during model fitting. A cell-cell neighborhood was calculated on this latent representation using the *sc.pp.neighbors* function, with *n_neighbors=int(0.25 * sqrt(adata.n_obs))*, and this cell-cell neighborhood was used to generate a 2-dimensional UMAP using the *sc.tl.umap* function with *init_pos="spectral"* (14). Cell clustering was performed using the leiden algorithm via the *sc.tl.leiden* function (15).

Differential gene expression testing was performed using the pyDEseq2 (16) and decoupler (17) python libraries, with cells pseudobulked by patient and cell type. Genes were filtered before testing using the *dc.pp.filter_by_prop* function with *min_prop=0.5* and *min_samples=int(0.33 * n_obs)*. Genes were considered differentially expressed if they had an absolute log-fold change of greater than 0.5 and an adjusted p-value of less than 0.05. Pathway analysis was performed on the results of differential gene expression tests using the *dc.mt.ulm* univariate linear model scoring function in decoupler against the PROGENy (Pathway RespOnsive GENes) pathway database (18). Over-representation analysis (ORA) of Gene Ontology (GO) categories was performed on differentially expressed genes using gProfiler (19).

Single-molecule spatial transcriptomics counts data generated on the 10X Genomics Xenium platform were analyzed using the methods described above, with minimal cell/gene filtration.

Visualizations of these data were generated using the squidpy (20) and spatialdata (21) python libraries.

**Histology and Immunohistochemistry.** Histology with hematoxylin and eosin staining was performed, and a pathologist confirmed disease for each PVOD sample. IHC staining was performed based on our established protocols (2, 5, 22). In brief, tissue samples were fixed overnight in 2% PFA, washed in 1X PBS, dehydrated in increasing gradation of ethanol (30%, 50%, 70%, 95% and 100%) and embedded in paraffin (Leica). 6-μm-thick cut sections were obtained; gross morphological changes were evaluated using hematoxylin and eosin staining. Subsequent sections were used for IHC, where antibody binding sites were exposed using HIER process, then incubated with 3% hydrogen peroxide for 10 min at room temperature to quench peroxidases activity and stained with primary antibody (Table 2) overnight at 4°C. Sections were then incubated with Alexa Fluor secondary antibodies (Table 3) to visualize the presence of relevant proteins. The slides were mounted using Vectashield Antifade Mounting Medium (H-1000, Vector Labs) and were imaged using Leica DMi8 confocal microscope.

**Supplemental Figure Legends**

**Figure E1.** Vascular morphological changes associated with PVOD. (A-D) Images of other H&E-stained tissues from unaffected (A-B) and PVOD (C-D) lungs, showing luminal occlusion and muscularization in PVOD lungs. (E) UMAP of embedded cells colored by unbiased Leiden based clustering. Scale bar: A-D=100µm.

**Figure E2.** Differential gene expression of selective EndMT gene markers. (A) Dot plot analysis of selective EndMT gene markers across ECs in cell atlas.

**Figure E3.** Expansion of systemic venous shunts in additional PVOD patients. (A-B) IHC staining for ACKR1, COL15A1and DAPI in PVOD tissue highlighting the presence of systemic veins and systemic venous shunts throughout the tissue, with colocalization of COL15A1 and ACKR1 in both venous shunts (A), airway systemic veins (B), and distal veins (C). Scale bars: 100µm.

**Figure E4.** Differential gene expression of ISR components in ECs and ferroptosis markers in all macrophages. (A-D) UMAPs of embedded ECs split by condition and colored by expression of *ATF4* (A), *EIF2AK2* (B), *GADD45A* (C) and *BRD4* (D). (E) Heat map of expression of marker genes for ferroptosis split by unaffected (left) and PVOD (right). (F) UMAP embedded by cell types. (G-I) UMAPs of embedded cells, split by conditions, and colored by gene expression of ferroptosis gene markers, *ACSL4* (G), *HMOX1* (H) and *TFRC* (I).

**Figure E5.** Cell type specific Increased TGF-ß pathway gene expression in PVOD. (A) UMAP of embedded ECs colored by cell subtypes and split by unaffected (left) and PVOD (right) status. (B-J) UMAP of embedded cell colored by expression of *SERPINE1* (B), *TGFBR1* (C), *TGFBR2* (D), *TGFBR3* (E), *BMP6* (F), *ACVRL1* (G), *ACVR1* (H), *ACVR1B* (I) and *ACVR2B* (J).

**Figure E6.** Spatially resolving expression of TGFβ and INHB family members across PVOD tissue. (A) Xenium spatial transcriptomics cluster assignments across PVOD tissue, showing only Mesenchymal, Endothelial, and Immune clusters. (B) Expression of *CDH5* (left) and *NOTCH3* (right) to localize endothelium and vascular smooth muscle/pericytes, respectively. (C) Expression of the activin/inhibin subunits INHBA and *INHBB*, as well as *MARCO*, a lung macrophage marker which co-localizes with *INHBA* expression. D,E) Expression of the TGFβ receptors *TGFBR1*, *TGFBR2*, and *TGFBR3* in (D) mesenchyme, and (E) endothelium. F,G) As in (D,E) for the TGFβ ligands *TGFB1*, *TGFB2*, and *TGFB3* in (F) mesenchyme, and (G) endothelium. (H,I) As in (D,E) for the expression of the activin receptors *ACVR1* and *ACVR1B* in (H) mesenchyme, and (I) endothelium.

**Figure E1**

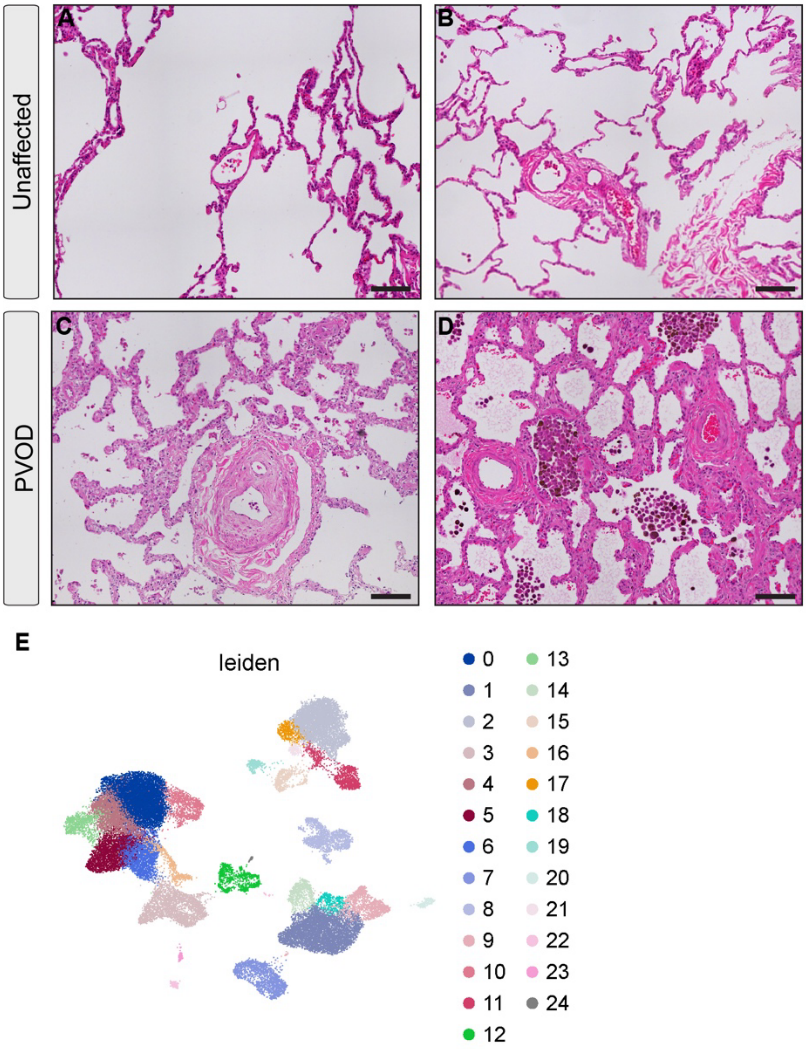
A
B
C
D
Unaffected
PVOD
E
leiden

**Figure E2.**

**A**

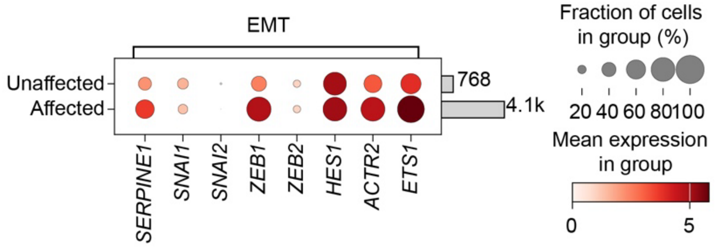

EMT
Unaffected
Affected
768
4.1k
SERPINE1
SNAI1
SNAI2
ZEB1
ZEB2
HES1
ACTR2
ETS1
Fraction of cells in group (%)
20 40 60 80100
Mean expression in group
0
5

Figure E3

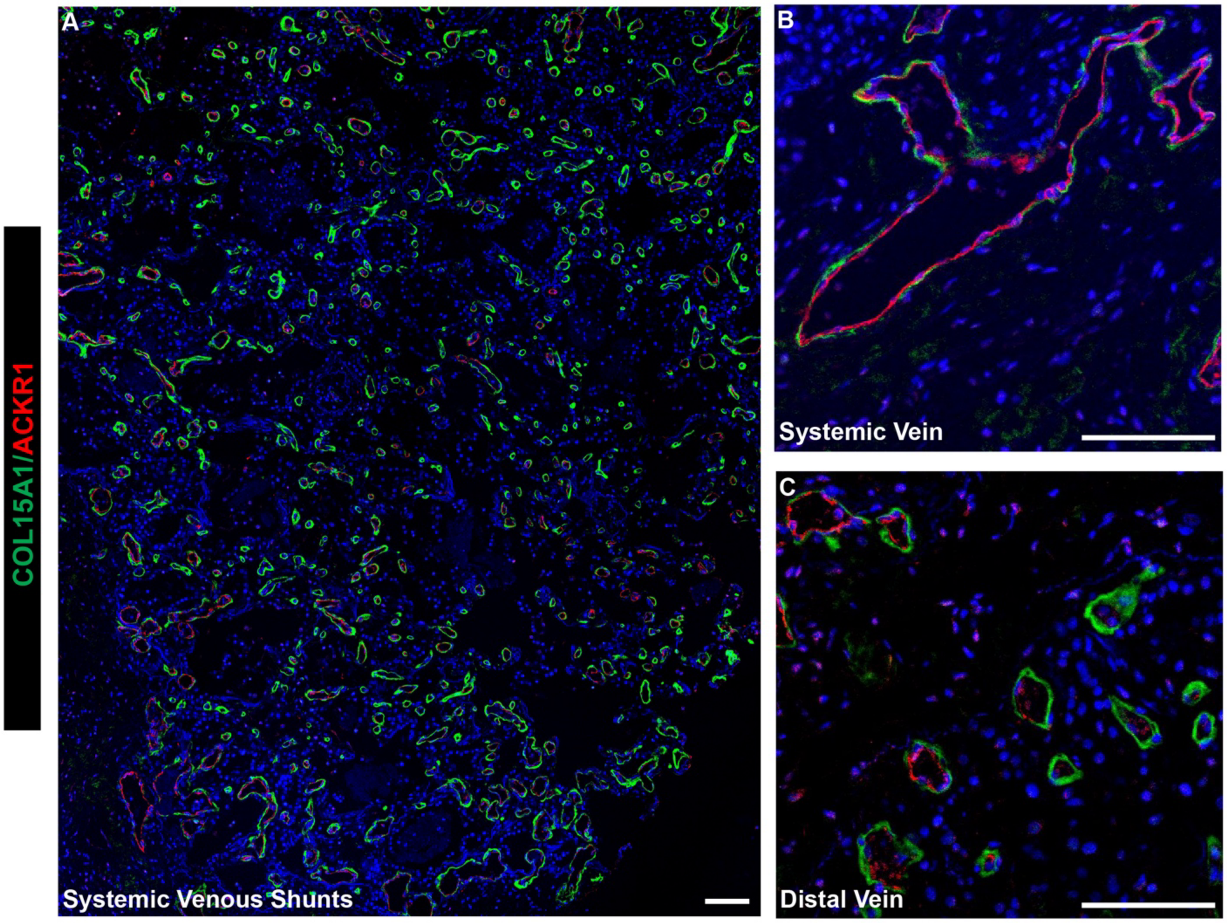

A
B
C
COL15A1/ACKR1
Systemic Venous Shunts
Systemic Vein
Distal Vein

### Figure E4

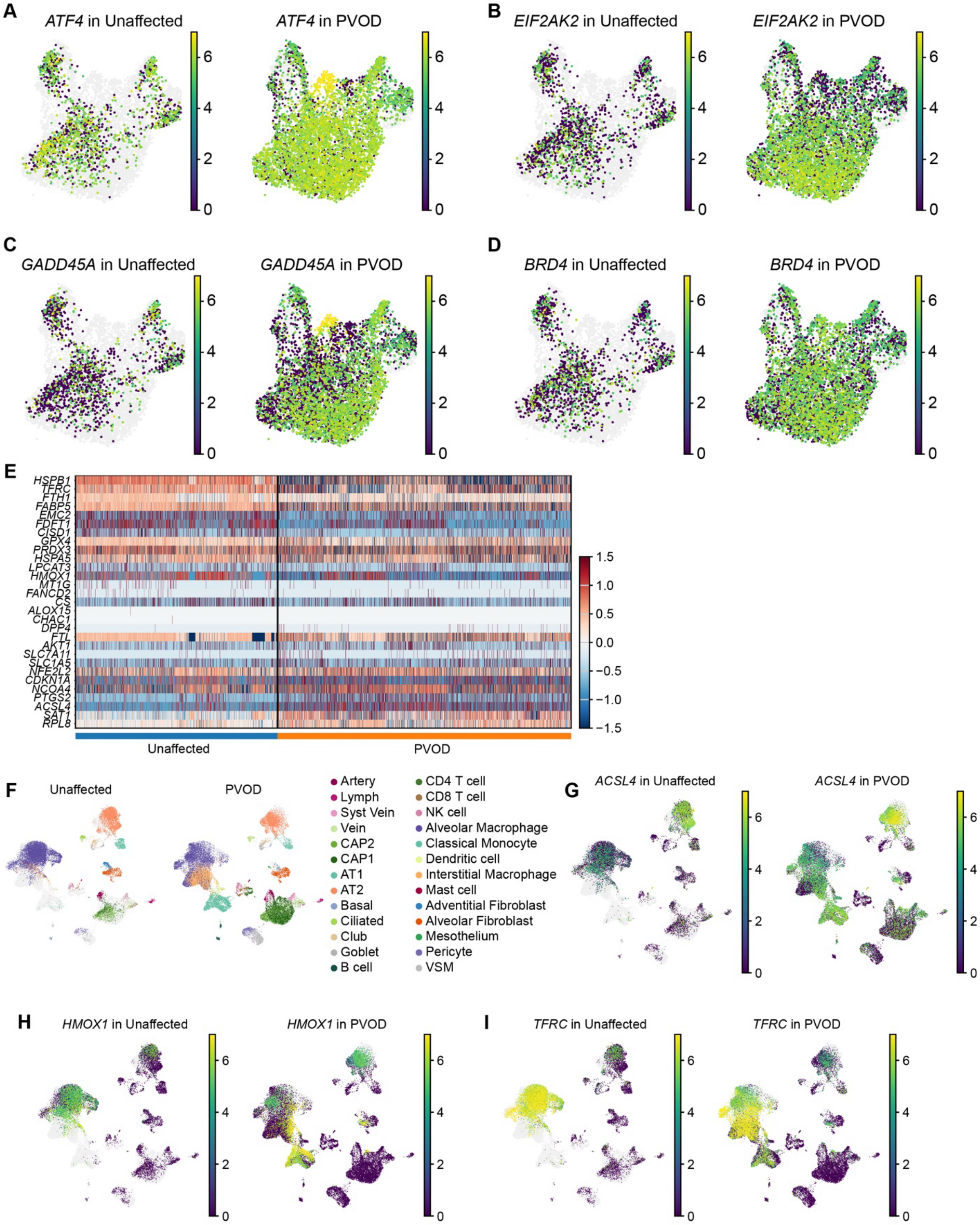
A
ATF4 in Unaffected
ATF4 in PVOD
B
EIF2AK2 in Unaffected
EIF2AK2 in PVOD
C
GADD45A in Unaffected
GADD45A in PVOD
D
BRD4 in Unaffected
BRD4 in PVOD
E
HSPB1
TFRC
FTH1
FABP5
EMC2
FDFT1
CISD1
GPX4
PRDX3
HSPA5
LPCAT3
HMOX1
MT1G
FANCD2
CS
ALOX15
CHAC1
DPP4
FTL
AKT1
SLC7A11
SLC1A5
NFE2L2
CDKN1A
NCOA4
PTGS2
ACSL4
SAT1
RPL8
Unaffected
PVOD
F
Unaffected
PVOD
Artery
Lymph
Syst Vein
Vein
CAP2
CAP1
AT1
AT2
Basal
Ciliated
Club
Goblet
B cell
CD4 T cell
CD8 T cell
NK cell
Alveolar Macrophage
Classical Monocyte
Dendritic cell
Interstitial Macrophage
Mast cell
Adventitial Fibroblast
Alveolar Fibroblast
Mesothelium
Pericyte
VSM
G
ACSL4 in Unaffected
ACSL4 in PVOD
H
HMOX1 in Unaffected
HMOX1 in PVOD
I
TFRC in Unaffected
TFRC in PVOD

**Figure E5.**

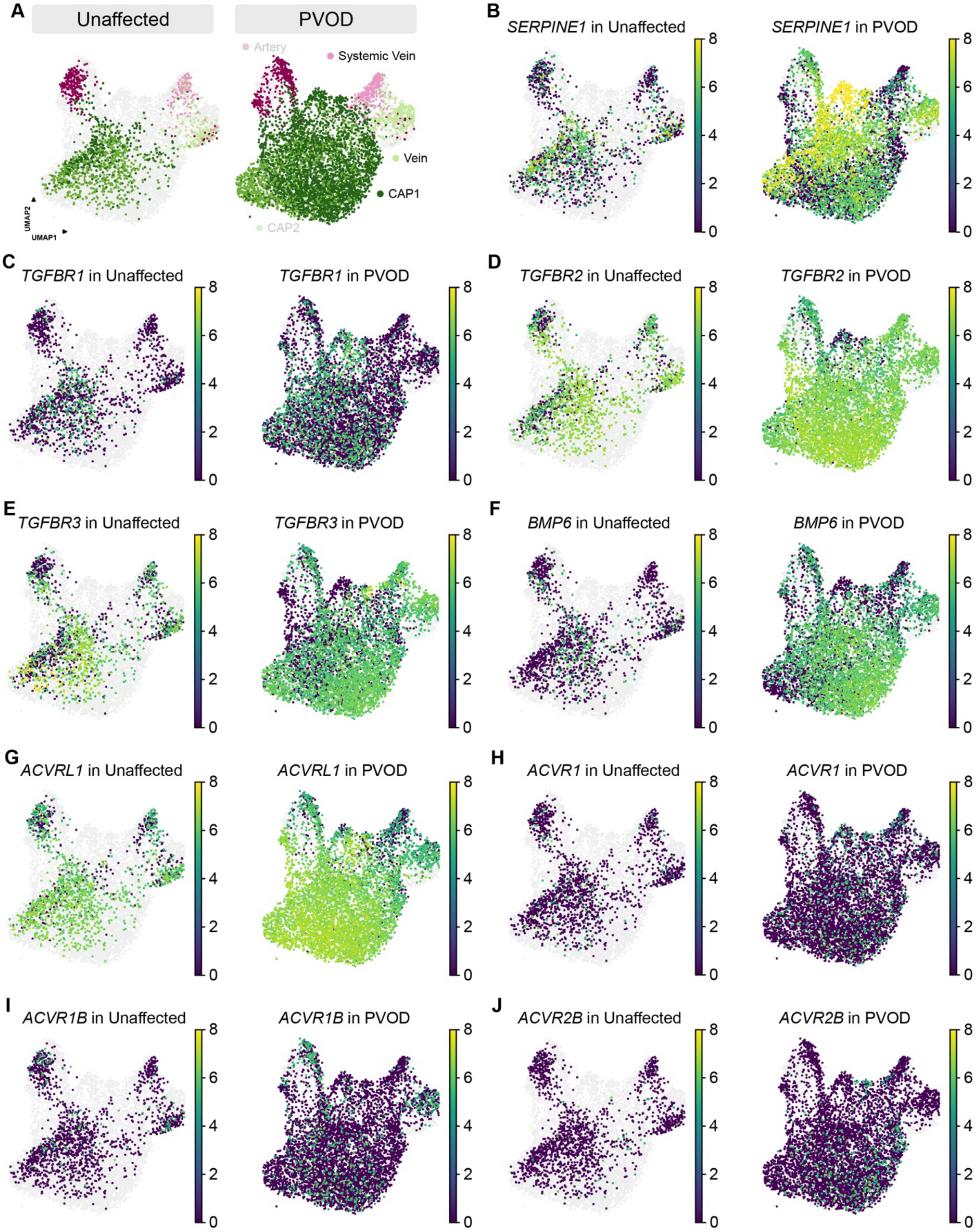

A
Unaffected
PVOD
Artery
Systemic Vein
Vein
CAP1
CAP2
UMAP2
UMAP1
B
SERPINE1 in Unaffected
SERPINE1 in PVOD
C
TGFBR1 in Unaffected
TGFBR1 in PVOD
D
TGFBR2 in Unaffected
TGFBR2 in PVOD
E
TGFBR3 in Unaffected
TGFBR3 in PVOD
F
BMP6 in Unaffected
BMP6 in PVOD
G
ACVRL1 in Unaffected
ACVRL1 in PVOD
H
ACVR1 in Unaffected
ACVR1 in PVOD
I
ACVR1B in Unaffected
ACVR1B in PVOD
J
ACVR2B in Unaffected
ACVR2B in PVOD
0
2
4
6
8

**Figure E6.**

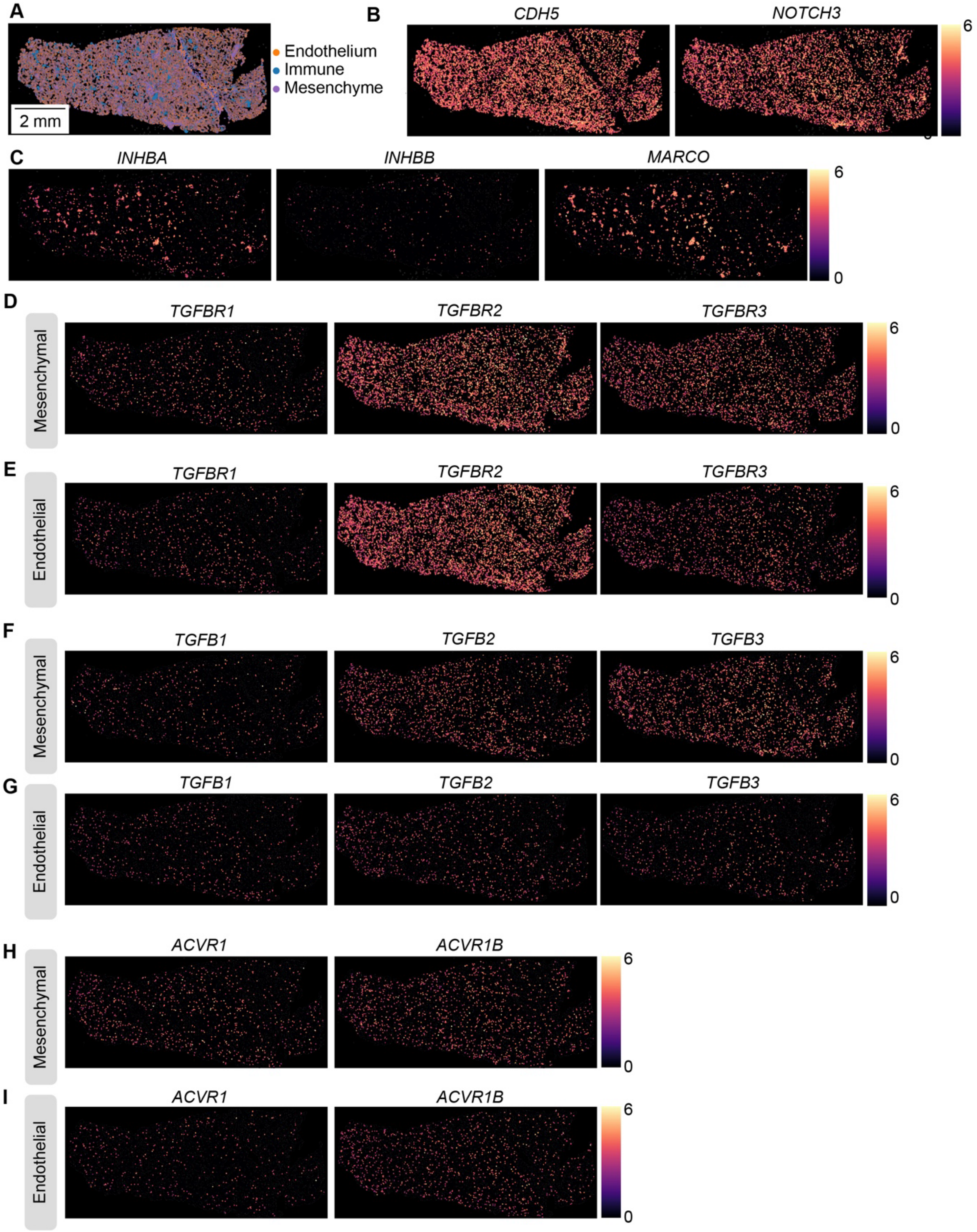
A
Endothelium
Immune
Mesenchyme
2 mm
B
CDH5
NOTCH3
6
C
INHBA
INHBB
MARCO
6
0
D
Mesenchymal
TGFBR1
TGFBR2
TGFBR3
6
0
E
Endothelial
TGFBR1
TGFBR2
TGFBR3
6
0
F
Mesenchymal
TGFB1
TGFB2
TGFB3
6
0
G
Endothelial
TGFB1
TGFB2
TGFB3
6
0
H
Mesenchymal
ACVR1
ACVR1B
6
0
I
Endothelial
ACVR1
ACVR1B
6
0
